# Spatially resolved transcriptomics of poplar reveals tissue organization across shoot-associated organs

**DOI:** 10.64898/2026.09.29.754462

**Authors:** Ziliang Luo, Anita Giabardo, Joshua C. Wood, Andrea Kohler, Chris Dardick, Chung-Jui Tsai, C. Robin Buell, Robert J. Schmitz

## Abstract

Poplar (*Populus* spp.) is a model for tree biology and a platform for engineering woody biomass, biofuels, biomaterials, and bioproducts. Many relevant traits depend on tissue position, developmental stage, and cell type, yet these spatial relationships are difficult to recover from bulk or single-cell transcriptomes. Here, we generated a spatial transcriptome atlas of *Populus tremula* × *P. alba* INRA 717-1B4 across the shoot apex, axillary bud, stem, and petiole. After quality control, the atlas retained 29,687 spatial spots from 45 tissue sections and detected 58,748 genes. Histology-guided clustering and marker analysis resolved meristematic, epidermal, cortical, vascular, and organ-specific domains. Cross-organ comparisons assessed whether published markers retained tissue-associated expression across different anatomical contexts and developmental stages, while *de novo* analysis identified additional domain-enriched candidates. As case studies of the utility of the atlas, we examined the emergence of trichome-associated programs in the shoot apex and adaxial-abaxial expression differences in petioles. A trichome identity score based on poplar markers from the single-cell shoot atlas peaked along the inferred meristem-to-primordium trajectory, revealing spatially localized expression of trichome-associated programs during early leaf development. Petiole expression differences were concentrated in the epidermis and cortex and involved polarity-associated, auxin-responsive, and cell-wall-remodeling genes, with distinct expression profiles across leaf positions. Together, these data provide a spatial reference for investigating tissue differentiation and developmental patterning in a transformable poplar genotype.

## Introduction

The genus *Populus* is a major model system for tree biology and an important platform for bioproduct and biomaterial research and production. As the first tree genome published, the *Populus trichocarpa* genome established a foundation for tree genomics and functional studies of secondary growth, wood formation, and environmental responses (Tuskan et al., 2006; Jansson and Douglas, 2007). Beyond its value as a model, poplar is an attractive feedstock for sustainable production of bioenergy, biomaterials, and bioproducts because of its rapid growth, vegetative propagation, and potential for genetic improvement and engineering (Buell et al., 2023; Sulis et al., 2025).

The hybrid genotype *Populus tremula* × *P. alba* INRA 717-1B4, referred to here as poplar 717, is widely used for functional genomics. Its haplotype-resolved genome and associated resources have strengthened poplar 717 as a reference genotype for allele-aware analysis and precision engineering (Zhou et al., 2023, 2025). Established genome-editing and base-editing workflows enable experimental tests of candidate regulators of development and biomass traits (Zhou et al., 2015; Triozzi et al., 2021; Li et al., 2021).

Single-cell and single-nucleus transcriptomics have begun to resolve cellular heterogeneity in *Populus*, including vascular, cambial, xylem, meristematic, and epidermal cell states (Chen et al., 2021; Conde et al., 2022; Li et al., 2023). A recent single-cell shoot atlas for poplar 717 provided a high-resolution reference for shoot cell types, epidermal heterogeneity, and trichome-associated developmental programs (Giabardo et al., 2026). However, single-cell approaches require isolation of protoplasts or nuclei and thereby lose the native tissue positions of the cells, limiting their ability to precisely identify where cell states and gene expression programs occur within intact organs.

Spatial transcriptomics addresses this gap by measuring gene expression in tissue sections, thereby preserving their spatial context. Since the original development of high-throughput spatial transcriptome mapping in tissue sections (Ståhl et al., 2016), plant spatial transcriptomics has expanded from proof-of-concept profiling to organ-scale and multiomic atlases. Early plant applications demonstrated spatially resolved transcriptome profiling in *Arabidopsis* inflorescences, *P. tremula* leaf buds, and Norway spruce (*Picea abies*) female cones (Giacomello et al., 2017). Subsequent work applied spatial approaches to metabolically and developmentally complex tissues, mapping integrated C4 and CAM photosynthesis within *Portulaca* leaves (Moreno-Villena et al., 2022), sucrose transport and storage programs in maize (*Zea mays*) kernels (Fu et al., 2023), and light-induced chlorenchyma programs during tomato (*Solanum lycopersicum*) shoot regeneration (Song et al., 2023). More recent studies have reached single-cell and cell-state resolution, resolving inflorescence meristem organization in the developing maize ear (Wang et al., 2024), endosperm developmental trajectories in a multiomic soybean (*Glycine max*) atlas (Zhang et al., 2025), a rare PRIMER cell state controlling localized immunity in *Arabidopsis* (Nobori et al., 2025), and spatiotemporal developmental programs governing conifer reproductive transitions in Norway spruce (Saarenpää et al., 2026). Together, these studies show that spatial transcriptomics is especially valuable in plant organs undergoing rapid changes in development, where gene expression is highly localized and follows a diverse set of cell developmental trajectories.

In *Populus*, published spatial and anatomically resolved transcriptomic resources have mainly focused on buds, wood formation, root regeneration, and leaf polarity. Spatially resolved profiling first captured developing and dormant *P. tremula* leaf buds (Giacomello et al., 2017). More recent studies integrated single-cell RNA sequencing with spatial transcriptome sequencing to map transcriptional domains during primary and secondary stem growth (Li et al., 2023), and combined complementary transcriptomic approaches to resolve xylem cell states and mechanical-stress responses during wood formation (Hsieh et al., 2025; Wei et al., 2026). These studies emphasize the value of anatomical information for interpreting cell identities and developmental relationships (Chen et al., 2024). Another study reconstructed *de novo* root regeneration trajectories in poplar cuttings (Lv et al., 2024). A single-nucleus and spatial atlas of poplar leaves further resolved adaxial-abaxial polarity and cuticle-deposition programs at cellular resolution (Li et al., 2026).

Building on this momentum and the recent single-cell atlas of poplar 717 (Giabardo et al., 2026), we developed a multi-organ spatial transcriptome atlas of poplar 717 across shoot development, including shoot apex, axillary bud, stem, and petiole. Because poplar 717 is widely used for transformation, genome editing, and functional genomics, a spatial reference in this genotype provides an anatomically informed atlas for interpreting gene expression programs, finding candidate regulators, and defining cell types and states in a matched genetic context.

Here, we used the multi-organ spatial transcriptome atlas of poplar 717 to address two questions about shoot development that require positional information. First, where do trichome-associated programs emerge during early leaf development in the shoot apex? Trichome density is an agronomically relevant trait in *Populus*, and locating these programs relative to the meristem and emerging leaf primordia provides spatial context for understanding trichome development. Second, how is adaxial-abaxial polarity transcriptionally organized in the petiole, the organ that sets leaf orientation and contributes to light interception in a tree canopy? Together, these analyses illustrate how the atlas connects developmental transcriptional programs to their anatomical positions within intact organs.

## Results

### A multi-organ spatial transcriptome atlas of poplar 717 shoot-associated tissues

We used the 10X Genomics Visium spatial platform to generate a spatial transcriptome atlas of poplar 717 shoot-associated organs, including the shoot apex, stem, axillary bud, and petiole (Fig. 1a). Histological sections captured distinct anatomical contexts for each organ, including longitudinal shoot apex tissue, stem cross sections, longitudinal axillary bud sections that included surrounding stem, petiole cross sections, and longitudinal sections of the petiole-stem junction (Fig. 1b).

After quality control, the atlas retained 29,687 spatial spots from 45 sections across the four organs. Across the atlas, 58,748 genes were detected from 64,160 annotated gene models in the haplotype-resolved poplar 717 genome. Sequencing depth varied across tissues and section types. Per-spot median gene and unique molecular identifier (UMI) counts were, respectively, 3,168 and 4,379 in the shoot apex, 7,353 and 13,569 in axillary bud, 995 and 1,359 in stem, 2,847 and 3,748 in petiole cross sections, and 3,441 and 4,846 in petiole longitudinal sections (Fig. 1c; Supplementary Table S1). Organelle-derived signal was low overall, with median combined mitochondrial and chloroplast fractions ranging from 0.19% in the axillary bud to 1.40% in the stem. In a joint uniform manifold approximation and projection (UMAP) analysis of all retained spatial transcriptomic spots across tissues, spots clustered largely by organ, with overlap among tissues that share anatomical components or similar section contexts (Fig. 1d).

**Figure 1.**
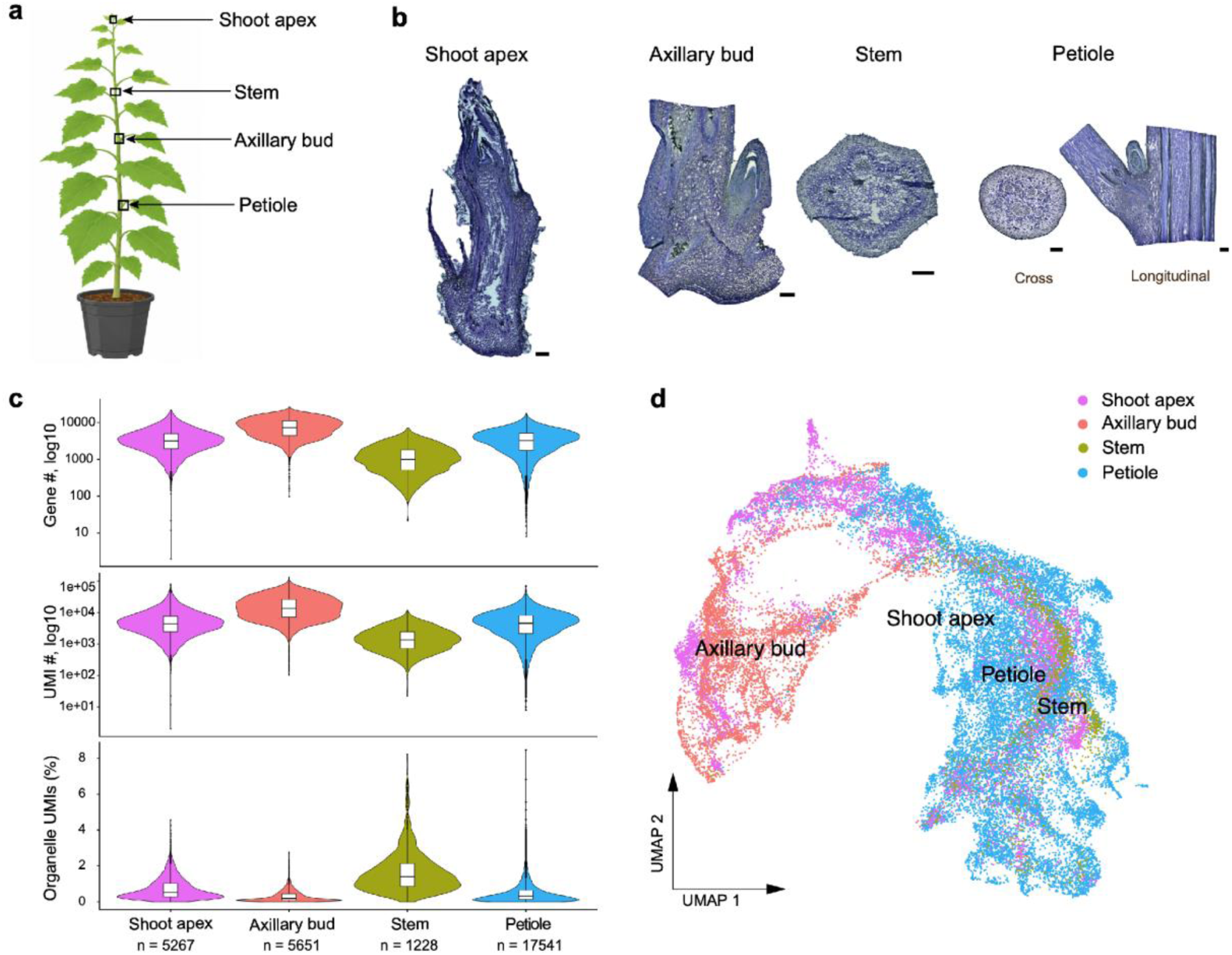
Generation and quality control of a multi-organ spatial transcriptome atlas of poplar 717 shoot-associated tissues. (**a**) Schematic showing sampled organs, including shoot apex, stem, axillary bud, and petiole. (**b**) Representative histological sections used for spatial transcriptome profiling. The atlas includes shoot apex, stem, axillary bud, and petiole sections, with petiole sampled in both cross and longitudinal orientations. Scale bars: 200 μm. (**c**) Quality-control distributions for detected gene number, UMI number, and mitochondrial plus chloroplast UMI percentage across shoot apex, axillary bud, stem, and petiole spots. Values are shown on log10 scales for gene and UMI counts. (**d**) Global embedding of spatial spots colored by sampled organ.

### Spatial clustering resolves anatomical domains across shoot apex, axillary bud, stem, and petiole

We next annotated spatial clusters within each organ based on histological position and domain-enriched marker gene expression. Because poplar 717 is an interspecific hybrid, mapping reads to its haplotype-resolved genome enabled separate quantification of *P. tremula* and *P. alba* alleles. Reciprocal syntelog pairs generally showed similar domain-level expression across all five spatial datasets (Supplementary Fig. S1). We therefore display the allele with the most specific and highest expression in the target domain. In the shoot apex, unsupervised transcriptome clustering resolved pith, cortex, epidermis, vasculature, leaf-primordium, and meristem-associated domains (Fig. 2a-c). Their positions matched the section anatomy: epidermal domains occurred at the surface, primordia formed emerging lateral organs, and vascular strands traversed internal ground tissues. Meristem-associated expression included *KNOTTED1-LIKE HOMEOBOX 2* (*KNAT2; PtXaAlbH.10G030700*), whose *Arabidopsis* homolog has been studied in shoot and floral development (Pautot et al., 2001). *PROLIFERATING CELL NUCLEAR ANTIGEN 2* (*PCNA2; PtXaTreH.01G204900*) and *CYCLIN A2;3* (*CYCA2;3, PtXaAlbH.01G152100*) provided evidence of proliferative states in young tissues (Imai et al., 2006; Strzalka et al., 2015). Epidermal annotation was supported by *PROTODERMAL FACTOR 1* (*PDF1; PtXaTreH.02G052300*), *3-KETOACYL-COA SYNTHASE 2* (*KCS2; PtXaAlbH.10G060600*), and *ECERIFERUM 5* (*CER5; PtXaTreH.01G212400*) homologs. These genes have established roles in protoderm identity, lipid elongation, and cuticular lipid export in *Arabidopsis* (Abe et al., 1999; Pighin et al., 2004; Joubès et al., 2008).

Stem marker expression followed the radial organization of the cross section (Fig. 2c). Pith-enriched expression of the established pith-associated marker *WRKY TRANSCRIPTION FACTOR 12* (*WRKY12*; *PtXaTreH.14G037200*) supported the pith annotation (Wang et al., 2010; Rao et al., 2019). *HYDROLASE* (*PtXaAlbH.18G083900*) and *PHOTOSYSTEM II SUBUNIT X* (*PSBXb*; *PtXaTreH.06G124900*) also showed pith-enriched expression in our stem sections (Fig. 2c); *PSBX* expression has previously been mapped to cortex and pith in poplar by spatial transcriptomics (Li et al., 2023). *PHLOEM INTERCALATED WITH XYLEM* (*PXY; PtXaTreH.01G106100*) supported cambium-associated annotation, consistent with functional evidence for *PXY* signaling in hybrid aspen (Etchells et al., 2015). Xylem-enriched expression of *4-COUMARATE:COENZYME A LIGASE 1* (*4CL1; PtXaAlbH.01G031600*) and *ENDO-BETA-MANNANASE 6* (*MAN6; PtXaAlbH.16G114900*) was consistent with their established roles in lignification and vessel development in *Populus* (Hu et al., 1998; Zhao et al., 2013; Zhou et al., 2015). *XYLEM CYSTEINE PEPTIDASE 1* (*XCP1*; *PtXaAlbH.04G160300*) and *XYLEM CYSTEINE PEPTIDASE 2* (*XCP2*; *PtXaAlbH.05G203100*) further supported the xylem annotation, consistent with published evidence of xylem expression in poplar (Nakaba et al., 2015; Gui et al., 2020; Fig. 2c). Across organs, phloem-associated domains expressed *SIEVE ELEMENT OCCLUSION-RELATED* (*SEOR*) and *PHLOEM PROTEIN 2-A* (*PP2-A*) homologs. *The SEOR* family includes the characterized *P. trichocarpa* locus *Potri.017G071000* (Kułak et al., 2025). The marker genes and corresponding poplar 717 loci are listed in Supplementary Table S2.

The axillary bud and petiole sections resolved additional organ-specific structures (Fig. 2a, b, Supplementary Fig. S2). In the axillary bud, spatial clustering separated bud primordium, bud procambium, bud scale, cortex, epidermis, phloem, vasculature, and xylem domains. Petiole cross sections contained five tissue-associated domains: epidermis, cortex, inner cortex, phloem, and xylem. Histology and marker expression supported these assignments (Supplementary Fig. S2; Supplementary Table S2). The phloem and xylem annotations reflect the predominant cell identity within each domain, although individual capture spots may include neighboring cell types. Across the four organs, the major spatial domains broadly corresponded to the manually outlined anatomical regions (Fig. 2a).

**Figure 2.**
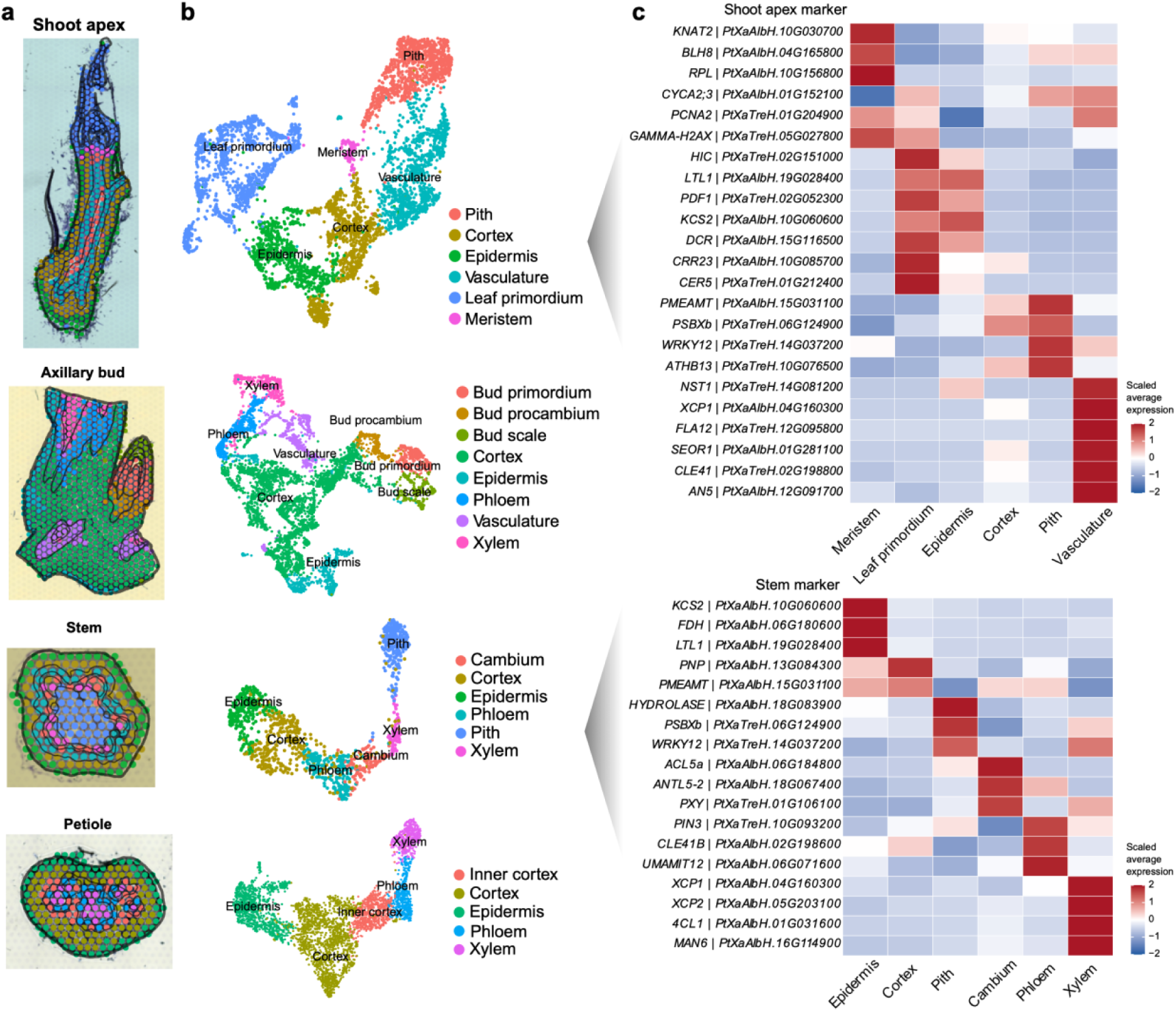
Spatial annotation of tissue domains across poplar shoot-associated organs. (**a**) Spatial spot maps overlaid on representative histological images for shoot apex, axillary bud, stem, and petiole sections. Spots are colored by annotated spatial domain. Black outlines mark tissue boundaries manually annotated from histological features. (**b**) UMAP embeddings of spatial spots for each profiled organ, colored by annotated spatial domain. (**c**) Heatmaps of scaled average expression for representative marker genes across annotated shoot apex and stem domains.

### Cross-organ marker consistency and *de novo* discovery extend the spatial annotation resource

We next examined whether known markers retained tissue-associated expression across organs differing in position and developmental stage (Fig. 3a). *KCS2* remained enriched in epidermal domains, *PHLOEM PROTEIN 2-A10-2* (*PP2-A10-2; PtXaAlbH.15G095300*) in phloem-associated domains, and *4CL1* in xylem-associated domains across the shoot apex, axillary bud, stem, and petiole. *XYLEM CYSTEINE PEPTIDASE 2* (*XCP2*) showed the same xylem-associated enrichment across organs, consistent with published evidence of xylem expression in poplar (Nakaba et al., 2015; Gui et al., 2020), and was more prominent in the axillary bud, stem, and petiole than in the shoot apex. *PXY* and the *WUSCHEL-RELATED HOMEOBOX 4* homolog *WOX4 (PtXaAlbH.14G016600)* marked cambium- and procambium-associated domains, consistent with the role of *PXY* signaling in vascular cambium activity in hybrid aspen (Etchells et al., 2015). *Populus WOX4*-like genes are required for normal cambial cell division activity and secondary growth (Kucukoglu et al., 2017). Spatial maps complemented this comparison by showing the distributions of *FIDDLEHEAD* (*FDH; PtXaTreH.06G179500*) and *PP2-A10-2* within intact organs (Fig. 3b). *PP2-A10-2* followed phloem-associated regions, whereas *FDH* showed broader expression in the shoot apex and axillary bud and clearer epidermal enrichment in the stem and petiole. These results identify tissue-associated expression patterns shared across anatomically and developmentally distinct organs, alongside markers whose distributions varied with organ context.

To identify novel cell-type-specific marker genes in poplar, we performed *de novo* marker discovery across the annotated tissues in the atlas. Candidates were ranked within each domain by how widely they were detected and how specific they were to that domain. We used the shoot apex to illustrate additional domain-enriched candidates across the meristem, emerging leaf primordia, and surrounding tissues (Fig. 3c, d; Supplementary Table S3). These candidates represented several functional classes and distinct spatial expression patterns. Along the shoot-apex surface, a peroxidase superfamily gene (*PtXaAlbH.04G010300*) and a GDSL-like lipase/acylhydrolase gene (*PtXaAlbH.11G053900*) were the two top-ranked epidermal candidates, detected in 99.6% and 68.0% of epidermal spots. In emerging leaf primordia, a bifunctional inhibitor/lipid-transfer protein gene (*PtXaTreH.16G108100*) and a chalcone-flavanone isomerase family gene (*PtXaTreH.19G056500*) were enriched. In internal regions, the methyltransferase-family gene *PtXaTreH.15G030300* and the NAD(P)-binding Rossmann-fold gene *PtXaTreH.14G011300* reached their strongest expression in the pith. In vascular-associated regions, a metallothionein 2A homolog (*PtXaTreH.09G023000*) and a leucine-rich repeat transmembrane protein kinase gene (*PtXaTreH.16G009200*) were enriched (Fig. 3c, d). Together, the cross-organ comparison and *de novo* discovery provide an *in situ* reference for assessing published markers and expand the set of candidate markers for tissues and spatial domains in poplar 717.

**Figure 3.**
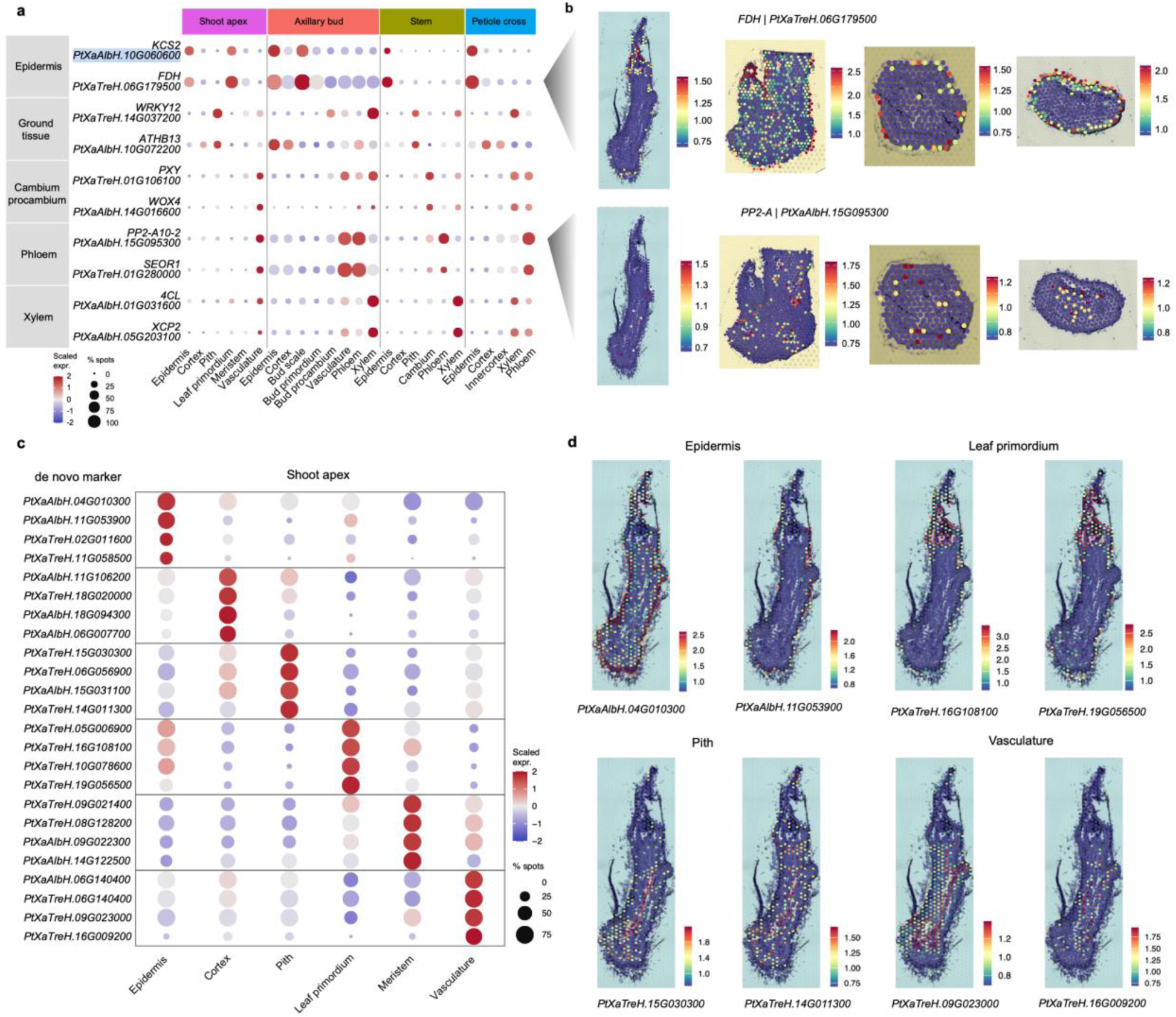
Cross-organ marker consistency and *de novo* domain-enriched candidates in the poplar shoot atlas. (**a**) Expression of published tissue-associated markers across annotated domains of the shoot apex, axillary bud, stem, and petiole. Color indicates scaled expression, and dot size indicates the percentage of expressing spots. (**b**) Spatial expression of *FDH* and *PP2-A* homologs across organ contexts. (**c**) Expression of *de novo* candidates across shoot-apex domains. (**d**) Spatial distributions of two of the top-ranked candidates for each of the epidermis, leaf primordium, pith and vasculature domains.

### Trichome-associated programs emerge during leaf primordium development

The poplar 717 single-cell shoot atlas identified trichome-associated epidermal states through expression of poplar trichome regulators, comparison between wild type and glabrous mutant bulk transcriptomes, and marker-promoter validation (Giabardo et al., 2026). The study interpreted epidermal clusters 14 and 39 from the scRNA-seq data as putative trichome initials and developing trichomes, respectively. In the single-cell reference, *MYB DOMAIN PROTEIN 38* (*MYB38*) was enriched in the putative initials, *SIAMESE-RELATED 1* (*SMR1*) was expressed in both states, and *MALONYLTRANSFERASE-LIKE 16* (*MATL16*) was enriched in the developing trichomes (Supplementary Fig. S3a). We used these three loci because they are trichome regulators with both mutant and marker-promoter evidence, and because all six of their poplar 717 alleles are highly expressed in the spatial shoot apex. We integrated this reference with our spatial data to locate trichome-associated programs within intact shoot-apex tissues. By using reference-based deconvolution, we mapped scRNA-seq clusters 14 and 39 predominantly to epidermal and leaf-primordium regions in the spatial atlas (Fig. 4a; Supplementary Fig. S4). A core trichome identity score, calculated as the per-spot mean of gene-wise standardized expression of the six poplar 717 alleles of *MYB38*, *SMR1* and *MATL16*, showed a corresponding spatial distribution (Fig. 4b). Together, these analyses localized trichome-associated transcriptional programs within developing shoot-apex tissues.

We identified expression-based subclusters within the meristem-to-leaf-primordium region and reconstructed a Monocle3 trajectory rooted in the meristem and extending toward lateral leaf primordia (Fig. 4c). Graph-based testing identified 4,390 genes associated with this trajectory (Fig. 4d; Supplementary Table S4). The six alleles of *MYB38*, *SMR1*, and *MATL16* showed elevated expression at later stages of the inferred trajectory. The core trichome score increased along pseudotime (Spearman ρ = 0.409; n = 1,426 spots), peaked within the leaf-primordium region, and declined toward the trajectory endpoint (Fig. 4e). Gene-specific scores revealed distinct expression peaks: *MYB38* and *SMR1* peaked before the combined score for *MATL16* (Supplementary Fig. S3b).

Co-expression analysis of the 1,426 meristem and leaf-primordium spots identified 17 modules (Supplementary Fig. S5a). Shoot apex-M4, Shoot apex-M7, and Shoot apex-M15 showed the strongest positive correlations with the core trichome score (Pearson r = 0.757, 0.539, and 0.444, respectively; Fig. 4f). Among these, Shoot apex-M4 also correlated positively with the predicted contributions of both trichome-associated single-cell clusters and with pseudotime (Fig. 4f; Supplementary Fig. S5b). Shoot apex-M4 activity was spatially concentrated within the developing shoot-apex region (Fig. 4g). The module was enriched for carbohydrate metabolism, cellulose and xylan biosynthesis, cell-wall organization, and responses to oxidative stress (Fig. 4h; Supplementary Fig. S5c; Supplementary Table S5). Its hubs included the proline-rich protein loci *PtXaTreH.04G133100* and *PtXaAlbH.04G132100*, together with expansin and GDSL-like lipase/acylhydrolase genes (Supplementary Fig. S5d; Supplementary Table S6). The proline-rich loci also varied along pseudotime and were expressed along the primordium and shoot-apex surface (Supplementary Fig. S6). These findings link the trichome-associated module to growth and cell-wall remodeling programs expressed at the developing shoot surface. Together, these analyses localized trichome-associated programs within developing leaf primordia and identified co-expression modules associated with their expression.

**Figure 4.**
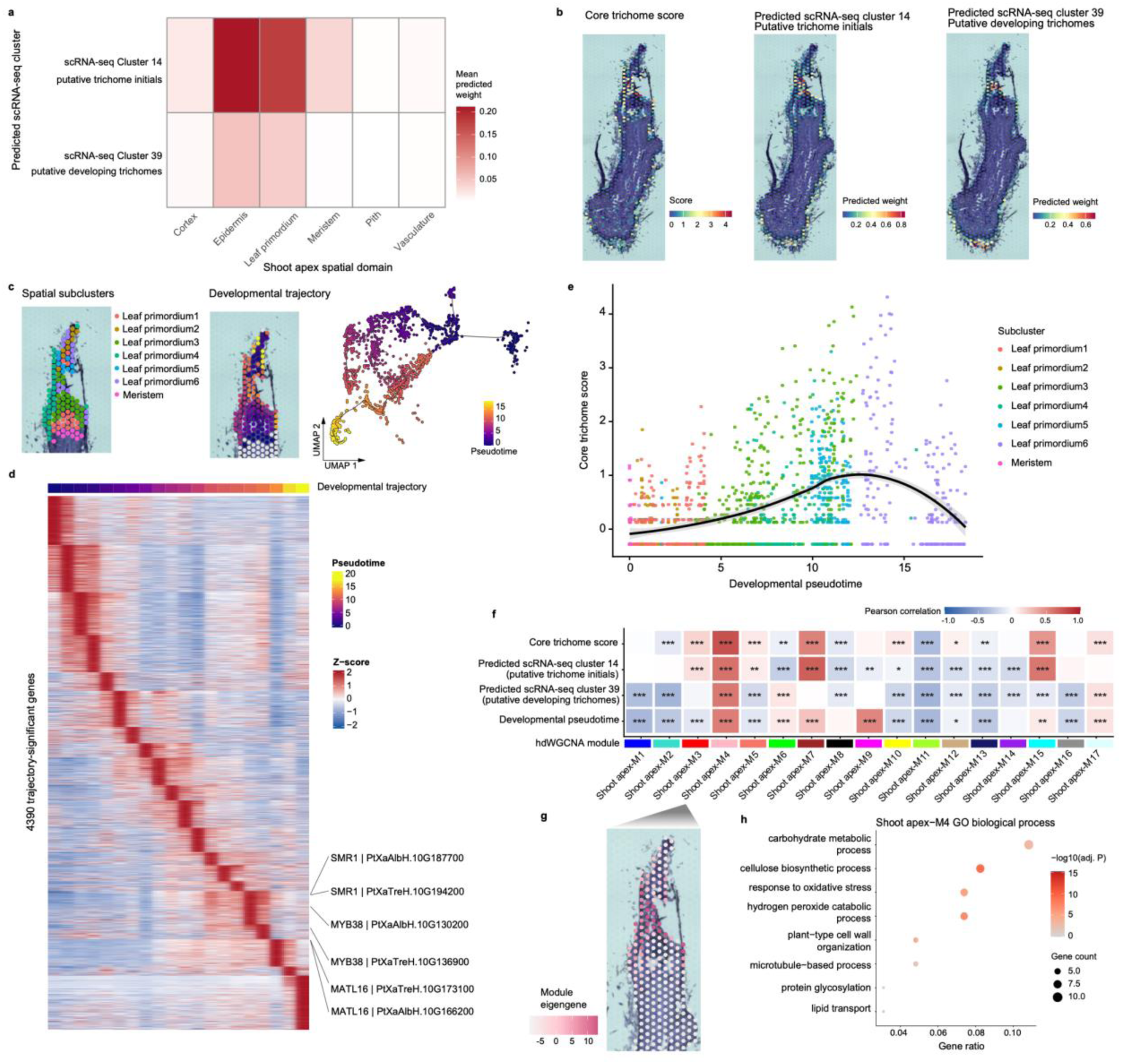
Spatial integration identifies trichome-associated programs during leaf primordium development. (**a**) Mean predicted contributions of single-cell reference clusters 14 and 39 across spatial domains, estimated by RCTD. These epidermal clusters were interpreted as putative trichome initials and developing trichomes, respectively, in Giabardo et al. (2026). (**b**) Spatial distribution of the core trichome score and predicted cluster 14 and 39 contributions in a representative shoot-apex section. The score is the mean standardized expression of six *MYB38*, *SMR1*, and *MATL16* alleles. (**c**) Seurat subclusters of the meristem-to-leaf-primordium region and the spatial and UMAP plots of the inferred Monocle3 trajectory. (**d**) Expression of 4,390 trajectory-significant genes along pseudotime, ordered by peak expression. Colors indicate gene-wise z-scores, capped at ±2. The positions of the six *MYB38*, *SMR1*, and *MATL16* alleles are indicated. (**e**) Core trichome score along inferred pseudotime. Each point represents a spatial spot, colored by developmental subcluster. The black curve shows the smoothed trend, and the shaded band indicates its 95% confidence interval. (**f**) Pearson correlations of 17 module eigengenes with the core score, predicted cluster 14 and 39 contributions, and pseudotime. Asterisks indicate Benjamini-Hochberg-adjusted P values (*P < 0.05, **P < 0.01, ***P < 0.001). The annotation bar indicates module colors. (**g**) Spatial distribution of Shoot apex-M4 eigengene expression. (**h**) Selected significantly enriched Gene Ontology biological processes for Shoot apex-M4. Dot size indicates gene count and color indicates −log10(adjusted P value).

### Adaxial-abaxial and developmental expression patterns in petiole epidermis and cortex

Petioles position leaf blades relative to the stem and contribute to light interception, mechanical support, and whole-shoot architecture. During poplar shoot development, changes in leaf and petiole morphology accompany vegetative development (Lawrence et al., 2021). Successive leaf positions provide a natural series for examining how petiole tissues acquire spatial asymmetry. We therefore used staged petiole cross sections to compare adaxial and abaxial transcriptional programs across tissue domains and developmental stages. Petiole sections from five leaf positions, L1, L2, L4, L5, and L15, were divided into adaxial and abaxial regions by anatomical orientation, resulting in 2,026 adaxial and 1,954 abaxial spots for side-specific analysis (Fig. 5a).

We identified 219 unique adaxial–abaxial DEGs in the epidermis or cortex across five leaf positions (Supplementary Table S7). These differences were concentrated in the epidermis and cortex (Fig. 5b). Abaxially enriched genes predominated at L1, L4 and L5, whereas few differences were detected at L2; at L15, most differences occurred in the cortex (Fig. 5c). Most of the DEGs (168 of 219) were detected in only one cell type and stage combination; the remaining 51 were detected in more than one combination (Supplementary Fig. S7). Genes with significant adaxial–abaxial differences were enriched for response to auxin, xyloglucan metabolism, cell-wall biogenesis and organization, calcium ion transport, ethylene-activated signaling, and response to oxidative stress (Fig. 5d; Supplementary Table S8). These terms point to epidermal and cortical programs involving hormone responses, wall remodeling, and growth-related signaling.

We next examined how the overall expression of these DEGs changed across petiole development. Three main expression groups were identified: genes with high expression at L1 that declined at later positions; genes with relatively high expression through L4 that declined at L5–L15; and genes with low early expression that increased from L4 onward (Supplementary Fig. S8a; Supplementary Table S9). The broad patterns were similar in the epidermis and cortex, with changes in several groups around L4–L5 petioles.

Among these developmentally varying DEGs, we identified three candidates with distinct adaxial–abaxial expression patterns and annotations related to polarity or growth: the *YABBY*-family locus *PtXaTreH.01G177100*, the *XTH6*-like locus *PtXaAlbH.07G007900* and the *SAUR*-like locus *PtXaTreH.09G107100* (Fig. 5e–k).

The *YABBY*-family gene *PtXaTreH.01G177100* had higher abaxial expression in the cortex across the sampled positions, with clear abaxial localization at L4 (Fig. 5e, f, i). Its best *Arabidopsis* hit is *YABBY2* (*YAB2*; AT1G08465), a member of the vegetative *YABBY* family studied in leaf polarity and development (Siegfried et al., 1999; Sarojam et al., 2010). The *XTH6*-like gene *PtXaAlbH.07G007900* increased in expression from L4 onward in the cortex, with higher abaxial expression at these positions (Fig. 5e). Spatial maps and spot-expression distributions at L4 showed this difference within the cortical domain (Fig. 5g, j). The *SAUR6*-like gene *PtXaTreH.09G107100* showed high early expression in the epidermis, lower expression at L4, an abaxial increase at L5 and no detected expression at L15 (Fig. 5e). Its abaxial enrichment at L1 was visible in the spatial map and spot-expression distribution (Fig. 5h, k). Additional epidermal examples showed different developmental profiles: the *EXPA5*-like gene *PtXaTreH.04G098000* peaked at L4, the *IAA4*-like gene *PtXaAlbH.05G168000* remained relatively high on the abaxial side through L5, and the *SAUR48*-like gene *PtXaTreH.06G108400* showed higher adaxial expression in most of the petioles (Supplementary Fig. S8b–e).

**Figure 5.**
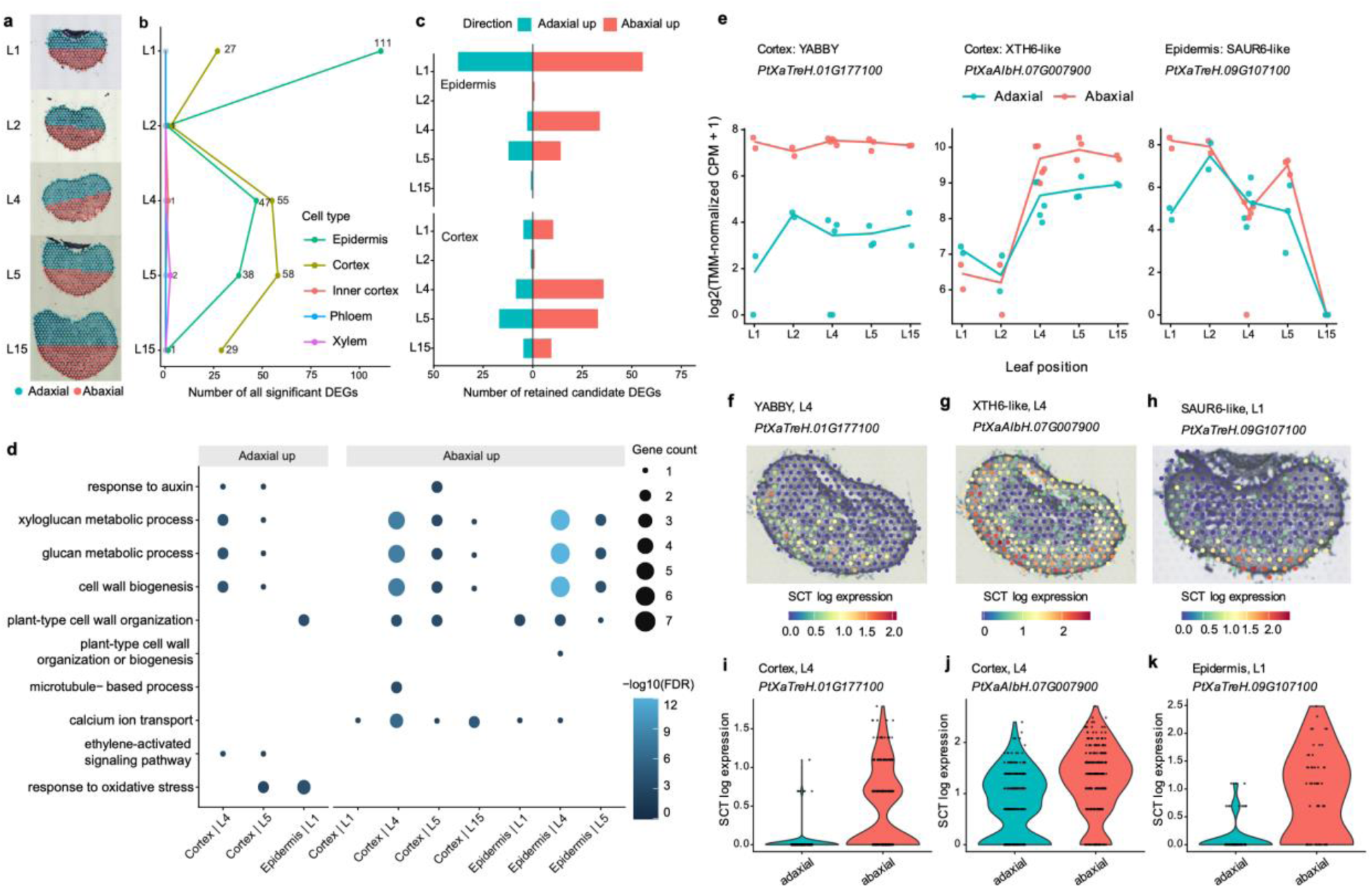
Adaxial–abaxial expression differences across petiole development. (**a**) Adaxial and abaxial regions in petiole cross sections at L1, L2, L4, L5 and L15. (**b**) Numbers of all significant DEGs by spatial domain and leaf position. (**c**) Numbers of retained candidate DEGs by enrichment direction and leaf position in the epidermis and cortex. Genes detected in multiple comparisons contribute to each corresponding count. (**d**) GO biological-process enrichment. (**e**) Adaxial and abaxial expression profiles of PtXaTreH.01G177100 and PtXaAlbH.07G007900 in the cortex, and PtXaTreH.09G107100 in the epidermis. Points represent individual sections; lines show log2-transformed equal-library mean CPM plus one. (**f**–**h**) Spatial expression of the same genes at L4, L4 and L1, respectively. (**i**–**k**) Corresponding side-specific distributions of SCT log-normalized expression within the indicated tissue. Points represent spatial spots pooled across sections and plants; distributions are descriptive.

## Discussion

This study provides a spatial transcriptome reference for shoot-associated organs in poplar 717, a genotype widely used for transformation, genome editing and biotechnology. Spatial transcriptomics retains positional information while measuring transcriptome-wide expression in tissue sections (Ståhl et al., 2016), and plant studies have increasingly used this approach to connect gene expression with anatomy, developmental patterning and tissue-specific regulation. Unlike earlier poplar spatial studies focused on a single tissue, this atlas combines multiple young shoot-associated organs from a transformable genotype with a matched single-cell reference.

Histology and published marker expression established anatomical-domain annotations, while cross-organ comparisons assessed marker consistency across plant positions and developmental stages. These comparisons identify markers with broadly retained tissue associations and those whose expression varies among organs. The *de novo* catalogue adds 988 marker entries representing 854 unique genes across 34 organ and section-specific tissue domains (Supplementary Table S3). Together, these spatial expression patterns and functional annotations provide a basis for selecting candidates for promoter testing and gene perturbation.

The shoot-apex analysis links trichome-associated transcriptional states to their positions within intact meristem and primordium tissues. Poplar trichome regulation is supported by functional studies of *MYB DOMAIN PROTEIN 186* (*MYB186*), *MYB DOMAIN PROTEIN 138* (*MYB138*), and *MYB38* (Plett et al., 2010; Bewg et al., 2022). The poplar 717 single-cell atlas associated epidermal subclusters epi_2 and epi_8 with leaf and shoot-apex trichome initials, respectively, and epi_11 with developing non-glandular trichomes (Giabardo et al., 2026). Their expression profiles distinguish initiation-associated *MYB38*, pan-trichome *SMR1*, and developing-trichome *MATL16* expression, providing a poplar-specific basis for interpreting the spatial data. Using these poplar markers, the spatial trichome score agreed with predicted trichome-cluster contributions and peaked along the meristem-to-leaf-primordium trajectory. The later peak of the developing-trichome signature relative to the initiation signature is consistent with progression of trichome-associated programs during early leaf development. Correlations with the core score identified Shoot apex-M4, Shoot apex-M7, and Shoot apex-M15 as associated gene modules. The spatial distribution and functional enrichment of Shoot apex-M4 further connect trichome-associated expression to cell-wall remodeling at the developing shoot surface.

Integrating the single-cell reference with spatial expression linked trichome-associated cell states to their anatomical positions and localized the associated co-expression module Shoot apex-M4 to developing leaf primordia and the shoot-apex surface. Within this module, the highest-ranked hubs were the paired PRP4-like loci PtXaTreH.04G133100 and PtXaAlbH.04G132100, whose expression also varied along pseudotime and localized to these developing surface regions (Supplementary Fig. S6; Supplementary Table S6). The module also contained the highly ranked EXPA8-like locus PtXaTreH.13G122000. Together with the module’s enrichment for cell-wall organization and biosynthesis, these candidates suggest that wall remodeling accompanies the emergence of trichome-associated programs. By linking single-cell identities and co-expression relationships with tissue position and developmental expression, this atlas identified a pool of candidates for testing their contributions to trichome initiation and development.

The staged petiole sections extended the atlas from organ initiation to lateral-organ maturation. Petioles coordinate vascular transport, mechanical support and leaf positioning, yet their spatial transcriptional organization is less well described than that of leaf blades or stems. In our data, adaxial-abaxial expression differences were concentrated mainly in the epidermis and cortex. This accords with the epidermal-growth-control hypothesis, which holds that in primary growth elongation and expansion are controlled by the epidermis (Savaldi-Goldstein et al., 2007; Kutschera and Niklas, 2007). This control has been linked to targeted cell wall loosening in the epidermis, and to hormone signaling between the epidermis and internal tissues (Savaldi-Goldstein et al., 2007; Kutschera and Niklas, 2007; Kutschera and Niklas, 2013). Many of the genes with adaxial-abaxial expression differences in the petiole fall into these two categories: cell wall remodeling, such as the *XTH6*-like locus *PtXaAlbH.07G007900* and the *EXPANSIN A5*-like locus *PtXaTreH.04G098000*, and hormone signaling, such as the *SAUR*-like locus *PtXaTreH.06G108400* and the *AUX/IAA*-family locus *PtXaAlbH.05G168000*. All these changes are likely governed by transcription factors that establish polarity in lateral organs. The strongest recurrent candidates were YABBY-family genes, consistent with the established role of YABBY transcription factors in abaxial identity in lateral organs (Siegfried et al., 1999; Bowman, 2000).

In *Arabidopsis*, far-red light perceived at the leaf tip promotes differential petiole growth through auxin transport and gibberellin signaling (Küpers et al., 2023). Our candidate list included loci annotated as homologs of *SAUR6*, *IAA4* and *XTH23*, whose *Arabidopsis* counterparts showed stronger transcriptional responses to this treatment in the abaxial petiole. These shared candidates are consistent with a possible contribution of auxin-responsive and cell-wall-related expression to petiole growth. The developmental profiles also showed that these candidate genes differed in the timing and persistence of expression across leaf positions. Their spatial distributions provide a basis for selecting genes for further study of petiole growth.

Several limitations should guide use of the atlas. First, Visium spots capture local tissue neighborhoods rather than individual cells, so spatial markers and deconvolution weights should be interpreted at the domain or local-neighborhood level. Second, sampling depth and replication differed across organs, making the atlas strongest for spatial annotation and candidate discovery rather than exhaustive quantitative comparison among all tissues. Third, the current petiole series captures developmental variation, but not controlled mechanical, light or gravity responses. Finally, marker conservation and candidate-gene prioritization require experimental validation before they can be treated as evidence of causal regulation.

The atlas supports functional studies in poplar 717 by placing candidate genes and promoters from bulk and single-cell studies within their anatomical context. Spatial expression, developmental trajectories and co-expression modules can guide selection of tissues and leaf positions for subsequent perturbation experiments. This application complements efforts to develop poplar for sustainable biofuels, biomaterials and bioproducts (Buell et al., 2023). Spatial expression, anatomical annotations, deconvolution weights, trajectories and module assignments are available through the BioPoplar Atlas website (https://bio-poplar-atlas.com/). Together with the poplar 717 single-cell resource, the atlas will allow users to examine candidate-gene expression across tissues and developmental stages.

## Methods

### Plant material and tissue collection

Clonally propagated female *P. tremula* × *P. alba* INRA 717-1B4 plants were grown as described by Giabardo et al. (2026). Spatial transcriptome profiling included the shoot apex, axillary buds, young stem at the 3^rd^ internode, and petioles. Stem cross sections were predominantly primary, with vascular bundles surrounding a central pith. Petioles were sampled at five leaf positions, L1, L2, L4, L5, and L15, using the leaf-position convention of Giabardo et al. (2026). Petiole sections came from the proximal region near the stem attachment.

Tissues were embedded, sectioned (12 µm thickness), stained with toluidine blue, imaged, and processed using the 10x Genomics Visium Spatial Gene Expression workflow. Sections comprised longitudinal shoot apex and axillary bud, stem cross sections, and petioles in both orientations. Histological images guided tissue alignment, spot selection, and spatial annotation.

### Spatial transcriptome preprocessing and quality control

Sequencing reads were aligned and counted against the haplotype-resolved poplar 717 genome assembly and v5 functional annotation (Zhou et al., 2023). The reference annotation contained 64,160 gene models. Spatial count matrices and histology images were imported into Seurat v5 (Stuart et al., 2019; Hao et al., 2024). Spots outside the tissue or the manually curated tissue masks were removed. For the cleaned shoot apex object, detached lower leaf spots were removed before final annotation and trajectory analysis. Quality-control metrics were summarized per tissue and section, including UMI counts, detected genes, and chloroplast, mitochondrial and combined organellar UMI fractions, excluding mitochondrial rRNA reads.

Spatial expression data were normalized using SCTransform (Hafemeister and Satija, 2019). Principal components were integrated across sections with Harmony, grouping by section identifier (Korsunsky et al., 2019). The first 30 Harmony dimensions were used for UMAP visualization, neighbor-graph construction, and clustering. Each organ and section orientation was analyzed separately because anatomy and cell-type composition differ.

### Spatial domain annotation and marker analysis

Spatial domains were annotated within each organ by combining histological position, spatial clustering, and marker-gene expression. Annotations were assigned separately for the shoot apex, axillary bud, stem, petiole cross sections, and petiole longitudinal sections. Candidate markers were curated from *Populus* studies, the poplar 717 single-cell atlas, and homolog-based evidence from *Arabidopsis* and other plants (Supplementary Table S2), and mapped to poplar 717 loci using functional annotation, homology and synteny. Marker heatmaps and dot plots informed the initial annotation. Cross-organ comparisons then tested whether domain-associated expression was retained; they are not independent validation. RCTD deconvolution added reference-based cell-state information for the shoot apex. In Figure 2a, tissue boundaries were outlined manually from histological features and overlaid on the domain maps.

*De novo* spatial markers were identified within each annotated tissue object using Seurat FindAllMarkers with the Wilcoxon rank-sum test. Positive markers were retained using Bonferroni-adjusted P < 0.01, average log2 fold change above 0.5 and detection in more than 25% of domain spots, after excluding curated known markers and organelle-encoded genes. Retained genes also required a domain-specificity log2 ratio above 0.5 between the mean expression in the target domain and the highest mean in any other domain of the same tissue, and were then ordered within each domain by detection fraction, mean expression, specificity and fold change. The atlas-wide marker table (Supplementary Table S3) reports the top-ranked genes of each shoot apex, axillary bud, stem and petiole cross-section domain, with gene identifiers, functional annotation, best *Arabidopsis* hit, log2 fold change, detection fractions, and adjusted P values. Because poplar 717 is an interspecific hybrid, most loci are represented by both a *P. tremula* and a *P. alba* allele. We compared the two alleles of each reciprocal syntelog pair, taken from the curated poplar 717 annotation and retained only where the assignment agreed in both directions (27,333 pairs), by their mean expression within every annotated spatial domain in which either allele was expressed (Supplementary Fig. S1). For each marker we show the allele with the highest, most domain-specific expression.

### Integration with the poplar 717 single-cell atlas

Shoot-apex single-cell profiles from the poplar 717 atlas served as the deconvolution reference for shoot-apex spatial transcriptomes (Giabardo et al., 2026). Reference and spatial samples came from the same genotype grown under matched conditions. Cell-state contributions to spatial spots were estimated using RCTD, implemented in spacexr, in multi-cell-type mode with a maximum of four cell types per spot, reflecting the mixed-cell composition of Visium spots (Cable et al., 2022). Predicted weights were summarized by annotated spatial domain. For the trichome analysis, epidermal reference clusters 14 and 39 were interpreted as associated with putative trichome initials and developing trichomes, respectively, following Giabardo et al. (2026). Their predicted contributions were mapped across the shoot apex and compared with the spatial trichome marker score.

### Shoot-apex trichome score, trajectory, and co-expression analysis

The core trichome identity score used the poplar 717 loci *MYB38* (*PtXaAlbH.10G130200* and *PtXaTreH.10G136900*), *SMR1* (*PtXaTreH.10G194200* and *PtXaAlbH.10G187700*), and *MATL16* (*PtXaAlbH.10G166200* and *PtXaTreH.10G173100*), following Giabardo et al. (2026). For each locus, SCT-normalized log-expression was standardized across spatial spots; the score was the mean of these six gene-wise z-scores. Stage-specific scores used the two *MYB38* alleles, the two *SMR1* alleles, or the two *MATL16* alleles. *LACCASE 3* (*LAC3*; *PtXaTreH.19G107500*) was excluded because it was represented by only 8 unique molecular identifiers across shoot-apex spots. The core score was projected onto tissue coordinates and the inferred meristem-to-leaf-primordium trajectory. Spearman correlations summarized its association with predicted single-cell cluster contributions and pseudotime.

Pseudotime was inferred from the shoot apex expression data with Monocle3 (Cao et al., 2019). The final cleaned shoot apex Seurat object was converted to a cell_data_set object, retaining the UMAP embedding and spatial-domain annotations. Cells were clustered, a principal graph was learned, and pseudotime was computed with the trajectory rooted in the meristem domain. Genes varying along the meristem-to-leaf-primordium branch were identified with graph_test and retained at q < 0.01 with positive Moran’s I. Only nuclear loci were kept for the trajectory-expression heatmap and Supplementary Table S4. For the full gene heatmap, mean SCT-normalized log expression was calculated in 20 equal-count pseudotime bins, standardized per gene across bins, and ordered by the bin of peak expression. The core trichome score was plotted against pseudotime with a LOESS fit (span = 0.7); stage-specific scores were summarized in ten equal-count pseudotime bins. The trajectory was inferred independently of the trichome scores.

Weighted gene co-expression analysis was performed with hdWGCNA on the 1,426 meristem and leaf-primordium spots used for trajectory analysis (Morabito et al., 2023). Genes detected in at least 5% of spots were retained. Metaspots were constructed within annotated domains and sections using 15 nearest neighbors, allowing a maximum overlap of eight spots and requiring at least 20 spots per group. A signed network used a soft-thresholding power chosen at a scale-free topology fit of 0.8 (fallback power 8). Modules were detected with a minimum size of 30 genes and merged at a dissimilarity threshold of 0.25. Module eigengenes were adjusted for section using Harmony. Pearson correlations were calculated between module eigengenes and the core trichome score, predicted contributions of reference clusters 14 and 39, and Monocle3 pseudotime, using spots with finite values for both variables. P values were adjusted across modules within each trait using the Benjamini-Hochberg procedure, and the same procedure was applied to the separate *MYB38*, *SMR1*, and *MATL16* scores. Module GO enrichment was tested with clusterProfiler using network genes as the background and GO terms containing 10-800 background genes (Yu et al., 2012). Hub genes were ranked by their correlations with the module eigengene.

### Petiole adaxial-abaxial analysis

Petiole cross sections were divided into equal adaxial and abaxial halves according to anatomical orientation. The initial analysis included 2,026 adaxial and 1,954 abaxial spots across L1, L2, L4, L5, and L15. We tested differential expression separately within each annotated domain and leaf position using the Wilcoxon rank-sum test in Seurat FindMarkers, requiring at least 15 spots per side, a minimum detection fraction of 0.1, and an absolute log2 fold-change threshold of 0.25. Significant comparisons had Bonferroni-adjusted P < 0.05. Positive and negative log2 fold changes indicated adaxial and abaxial enrichment, respectively. Spatial prioritization excluded tissue-identity markers, required a significant, directionally consistent correlation with the spatial side score (Benjamini-Hochberg-adjusted P < 0.05), and required the same side enrichment in at least two sections. Candidate and developmental analyses then focused on the epidermis and cortex. GO biological-process enrichment of the initial significant DEGs was tested separately by tissue, leaf position, and enrichment direction using clusterProfiler, with GO-annotated genes in the spatial object as the background and term sizes of 10-500 genes (Yu et al., 2012). P values were adjusted across terms within each gene set using the Benjamini-Hochberg procedure.

Developmental expression analysis used the union of 219 prioritized adaxial–abaxial DEGs, measuring each candidate in both the epidermis and cortex regardless of the tissue in which it was originally significant. Raw counts were summed by tissue, parent library and side, pooling serial sections within each library. TMM normalization was performed jointly across these samples (Robinson and Oshlack, 2010). For overall developmental profiles, normalized CPM values were averaged equally across sides within each library and then equally across libraries at each leaf position. Gene–tissue profiles were retained if they reached at least 10 CPM, spanned at least one log2(CPM + 1) unit, and had at least one position with detection in at least five spots and 10% of tissue spots in every available library. Each five-position log2(CPM + 1) profile was standardized separately before clustering.

The gene–tissue profiles were clustered jointly using Mfuzz (Kumar and Futschik, 2007), with an estimated fuzzifier of 2.182. Solutions with two to six clusters were evaluated using 30 random starts per cluster number, retaining the lowest-objective fit. Stability was assessed with 20 random subsets of 80% of genes, keeping both tissue profiles per sampled gene, and adjusted Rand indices were computed after predicting assignments for all profiles. Silhouette widths, Xie–Beni indices and centroid distances were also examined, and three clusters were used. Profiles were assigned to their highest-membership cluster, with membership ≥0.5 defining core profiles.

For side-specific candidate plots, section counts were divided by the section’s total gene-count depth and the corresponding parent tissue–side TMM normalization factor to obtain CPM. Points show individual sections, and lines show log2-transformed equal-library mean CPM plus one. Spatial maps and violin plots show SCT log-normalized spot expression. For the Figure 5 violins, spots from the indicated tissue and leaf position were pooled across sections and plants; the distributions are descriptive. Plant-specific distributions were checked separately; pooled spots were not treated as independent biological replicates. L4 and L5 each included samples from two different plants. Serial sections were treated as technical subsamples; different library identifiers at other positions were not assumed to represent different plants. Candidates were chosen on expression magnitude, spatial detection, consistency across the two plants at L4/L5, and gene annotation. An additional conservative screen required an original fold change >4, an enriched-to-other (CPM + 1) ratio >4 in every available library, and detection in at least five spots and 10% of enriched-side spots in each library. *XTH6*-like was retained as a well-detected illustrative candidate under the original DEG criteria rather than this additional screen. Leaf positions were displayed in anatomical order, without treating their spacing as elapsed time.

## Data Availability Statement

Raw sequencing data have been deposited in the NCBI BioProject database under accession number PRJNA1531574. All original codes are deposited at GitHub (https://github.com/Kennyluo4/Biopoplar_spatial_transcriptome_analysis). Spatial coordinates, domain annotations, marker tables, module assignments, deconvolution weights, and example gene-expression maps are accessible through the BioPoplar Atlas website (https://bio-poplar-atlas.com/) and associated data repositories. Analysis code is available through public repositories.

## Acknowledgements

The authors acknowledge Gilles Pilate (Institut National de la Recherche Agronomique, France) for providing poplar clone INRA 717-1B4. This material is based upon work supported by the U.S. Department of Energy, Office of Science, Office of Biological and Environmental Research program under Award Number DE-SC0023338.

## Author contributions

RJS, CD, CJT and CRB designed the research. ZL, JCW and AG performed experiments and analyzed data. ZL, JCW, AG, AK, CD, CJT, CRB and RJS wrote the manuscript. All coauthors approved the manuscript.

---

**Supplementary Figure S1.**
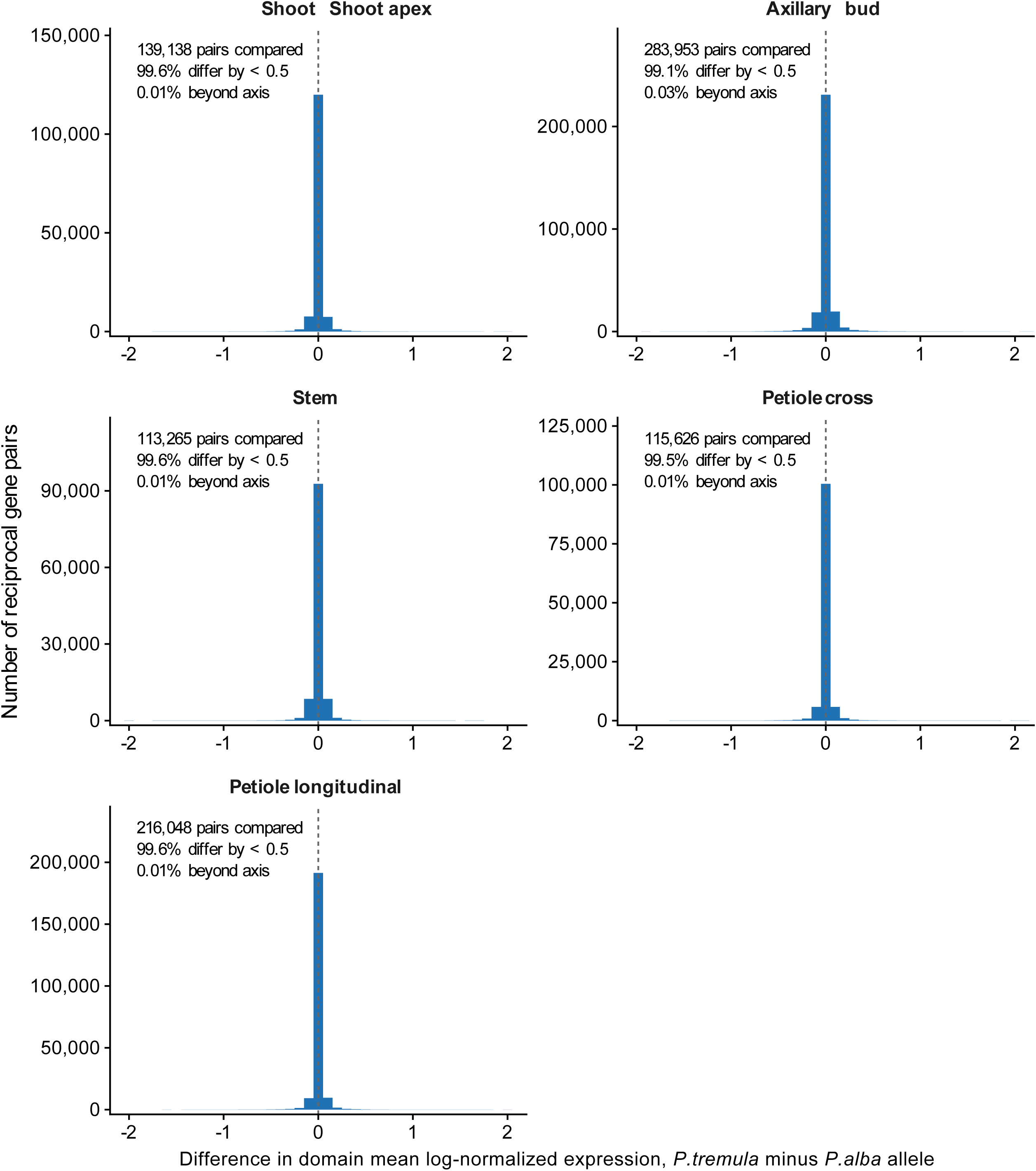
Domain-level expression of the *P. tremula* and *P. alba* alleles of reciprocal syntelog pairs differs little across the spatial atlas.

**Supplementary Figure S2.**
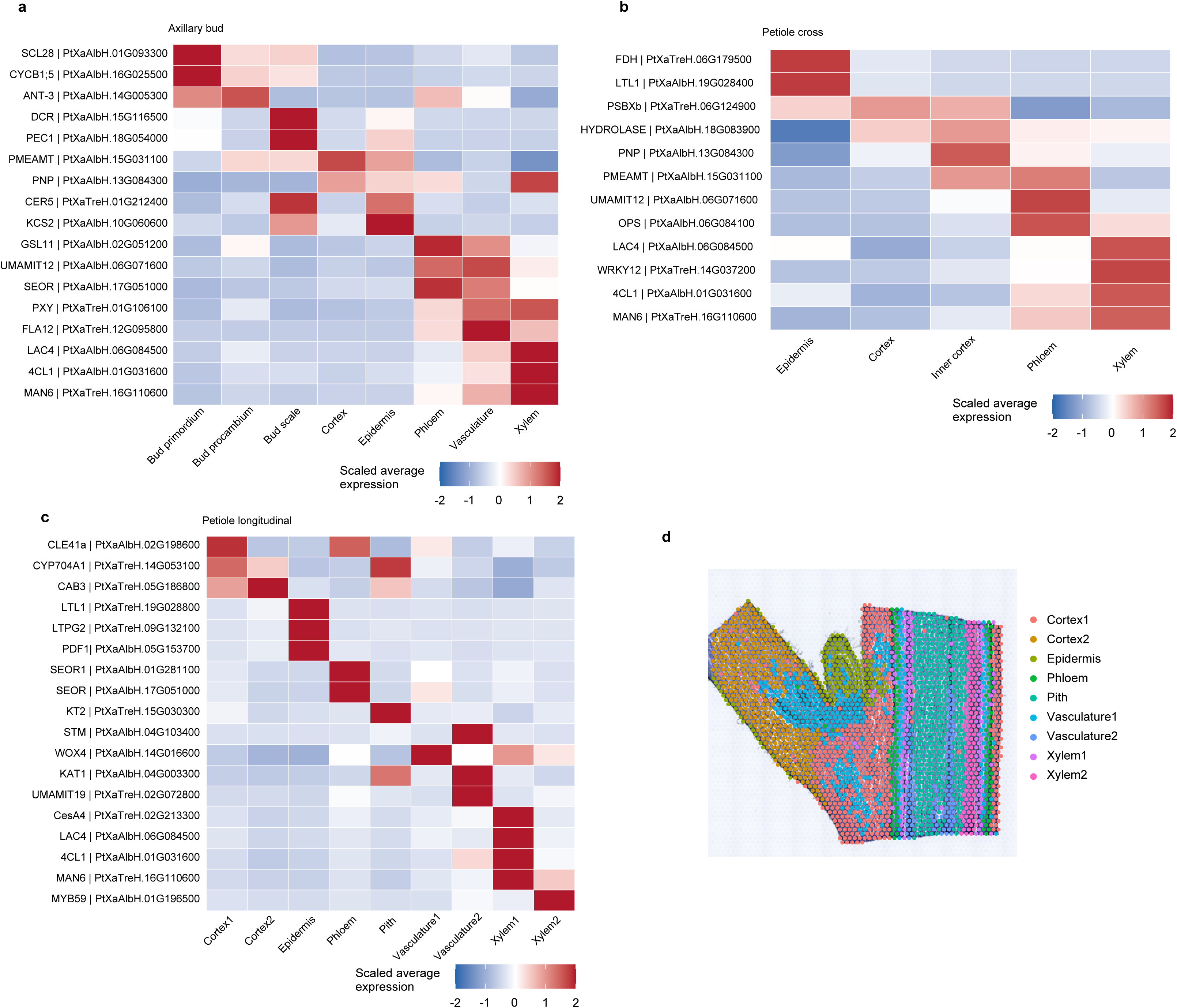
Spatial-domain annotation. Marker expression in axillary bud (a), petiole cross sections (b), and longitudinal sections (c). Colors indicate scaled mean expression. (d) Annotated domains in a petiole longitudinal section.

**Supplementary Figure S3.**
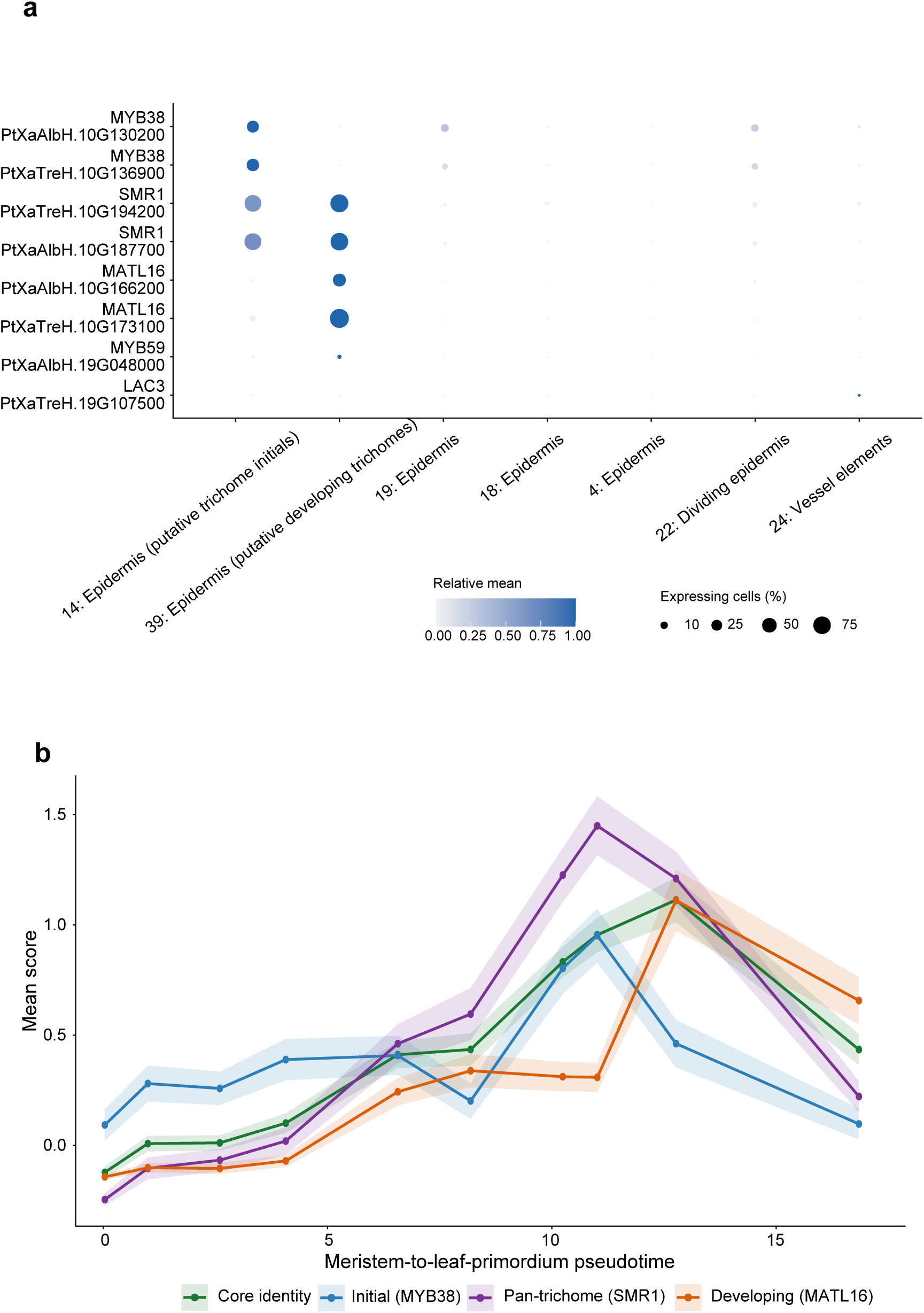
Trichome-associated markers and developmental scores. (a) Marker expression in selected single-cell reference clusters (Giabardo et al., 2026). Dot size indicates expressing-cell percentage; color indicates relative mean expression. (b) Trichome scores along pseudotime, shown as means ± standard errors.

**Supplementary Figure S4.**
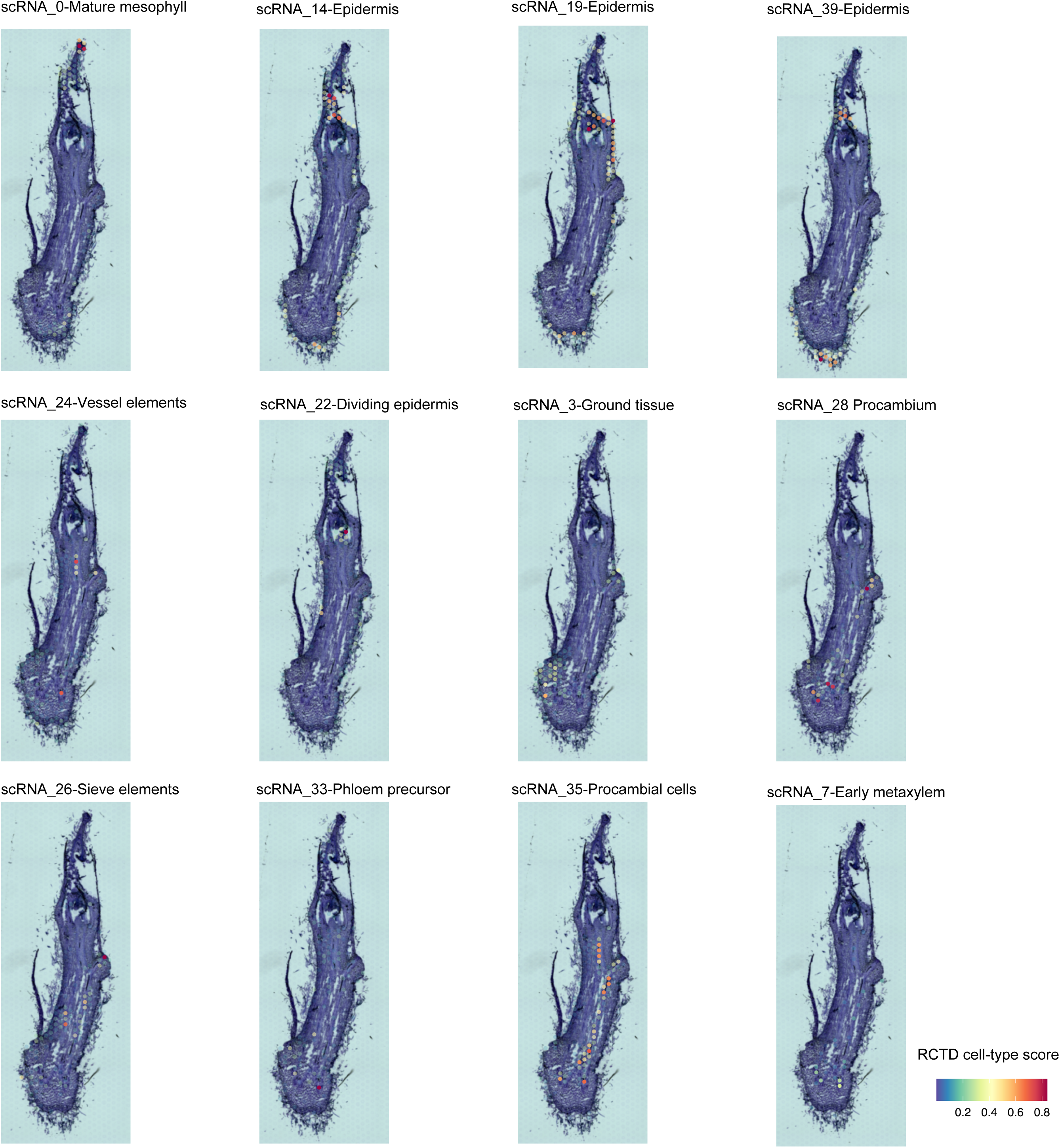
RCTD mapping of reference-cell states. Spatial distributions of predicted single-cell cluster contributions in the shoot apex.

**Supplementary Figure S5.**
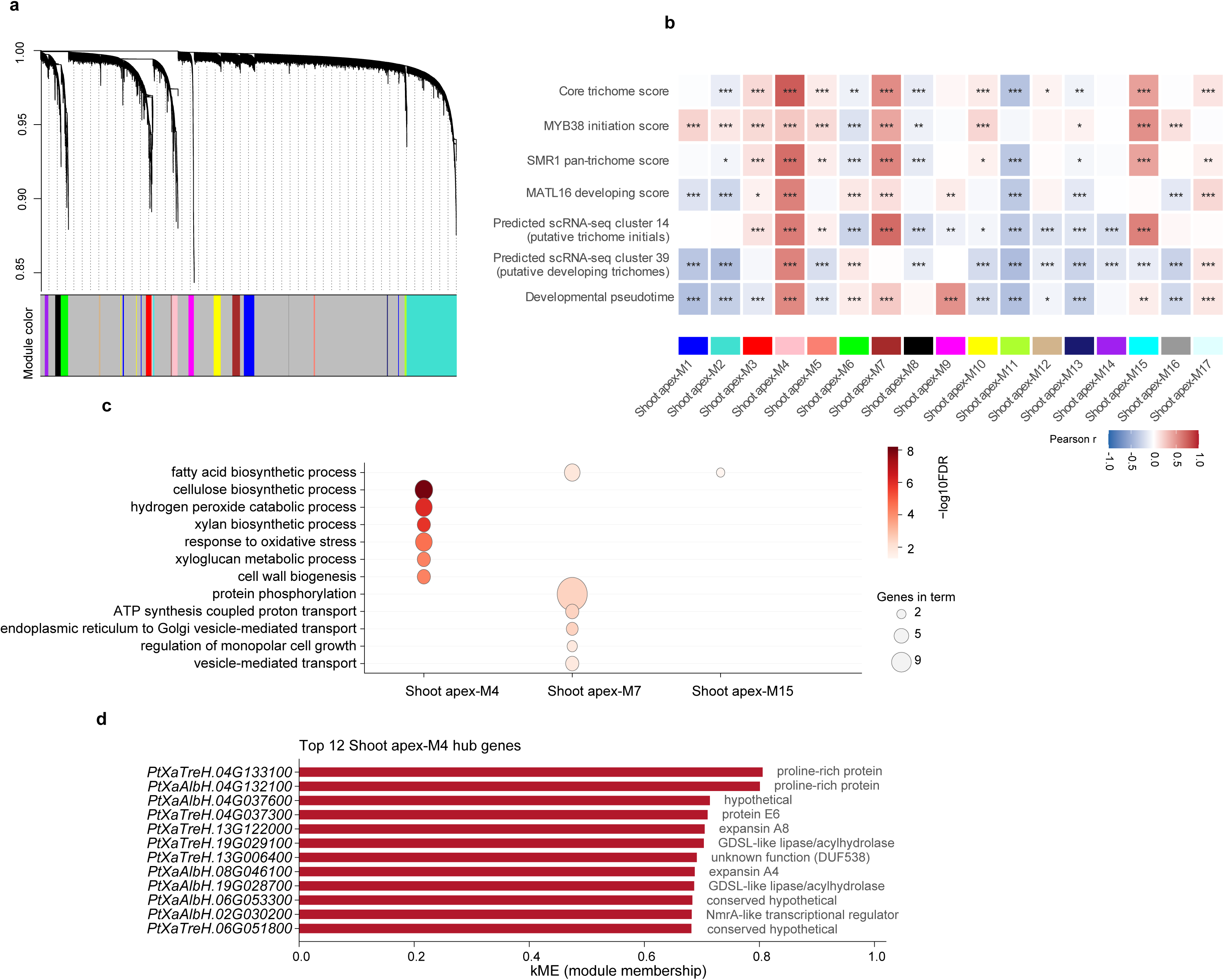
Trichome-associated co-expression modules. (a) Gene dendrogram and module assignments. (b) Module correlations with trichome scores, predicted cluster contributions, and pseudotime. Asterisks indicate adjusted P < 0.05 (*), < 0.01 (**), and < 0.001 (***). (c) GO enrichment of modules M4, M7, and M15; dot size indicates gene count and color indicates −log10(FDR). (d) Top 12 M4 hub genes ranked by kME.

**Supplementary Figure S6.**
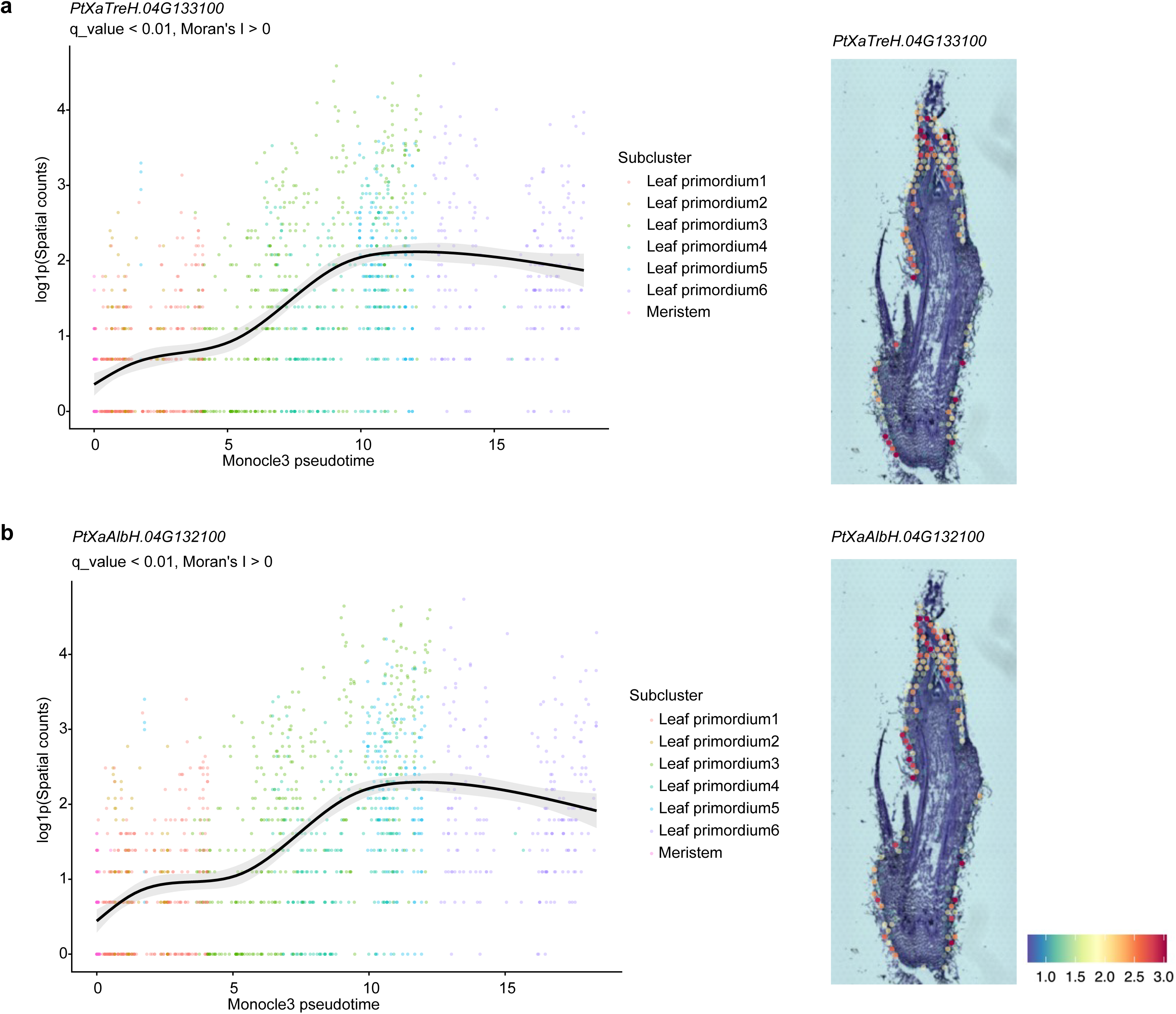
Expression of proline-rich protein genes. Developmental and spatial expression of *PtXaTreH.04G133100* (a) and *PtXaAlbH.04G132100* (b). Points represent spots colored by subcluster; curves show expression trends.

**Supplementary Figure S7.**
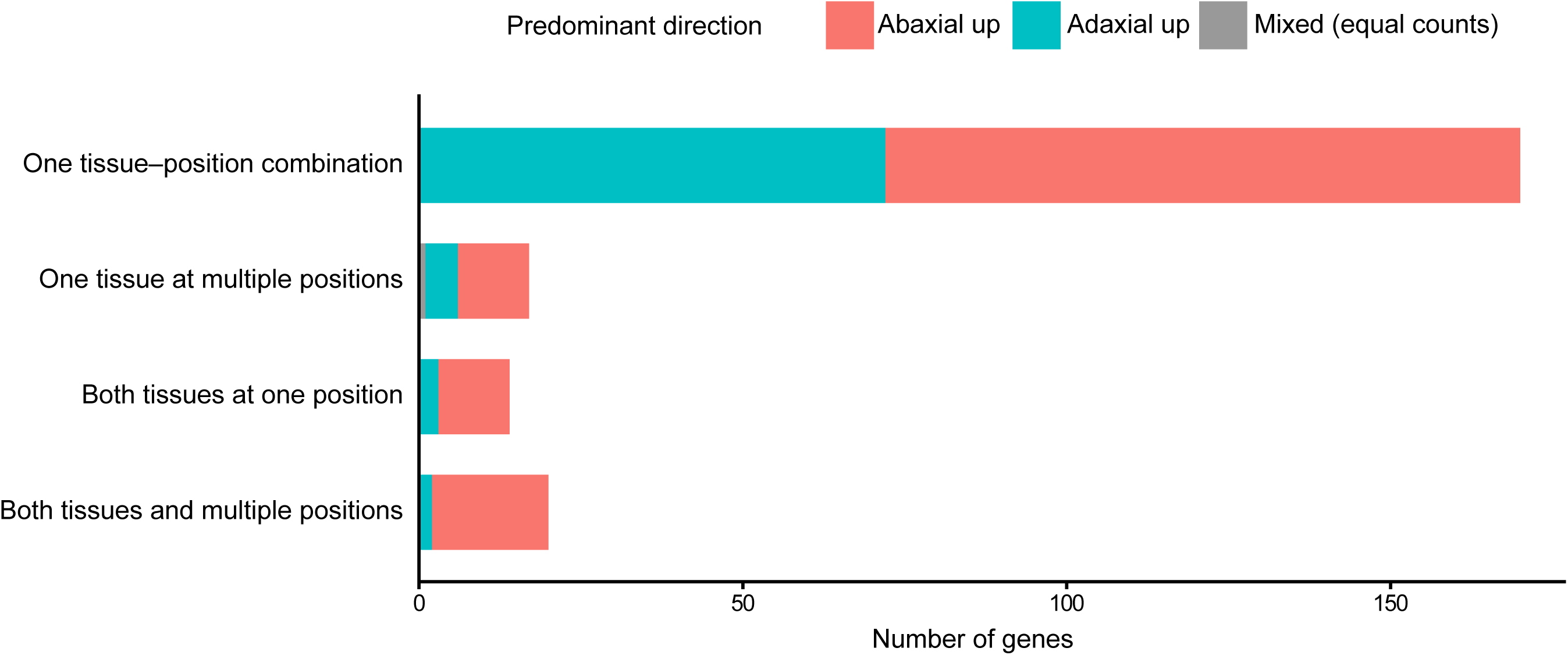
Recurrence of petiole adaxial–abaxial DEGs. Distribution of 219 genes across tissues and leaf positions. Colors indicate the predominant direction of enrichment; gray indicates equal recurrence on both sides.

**Supplementary Figure S8.**
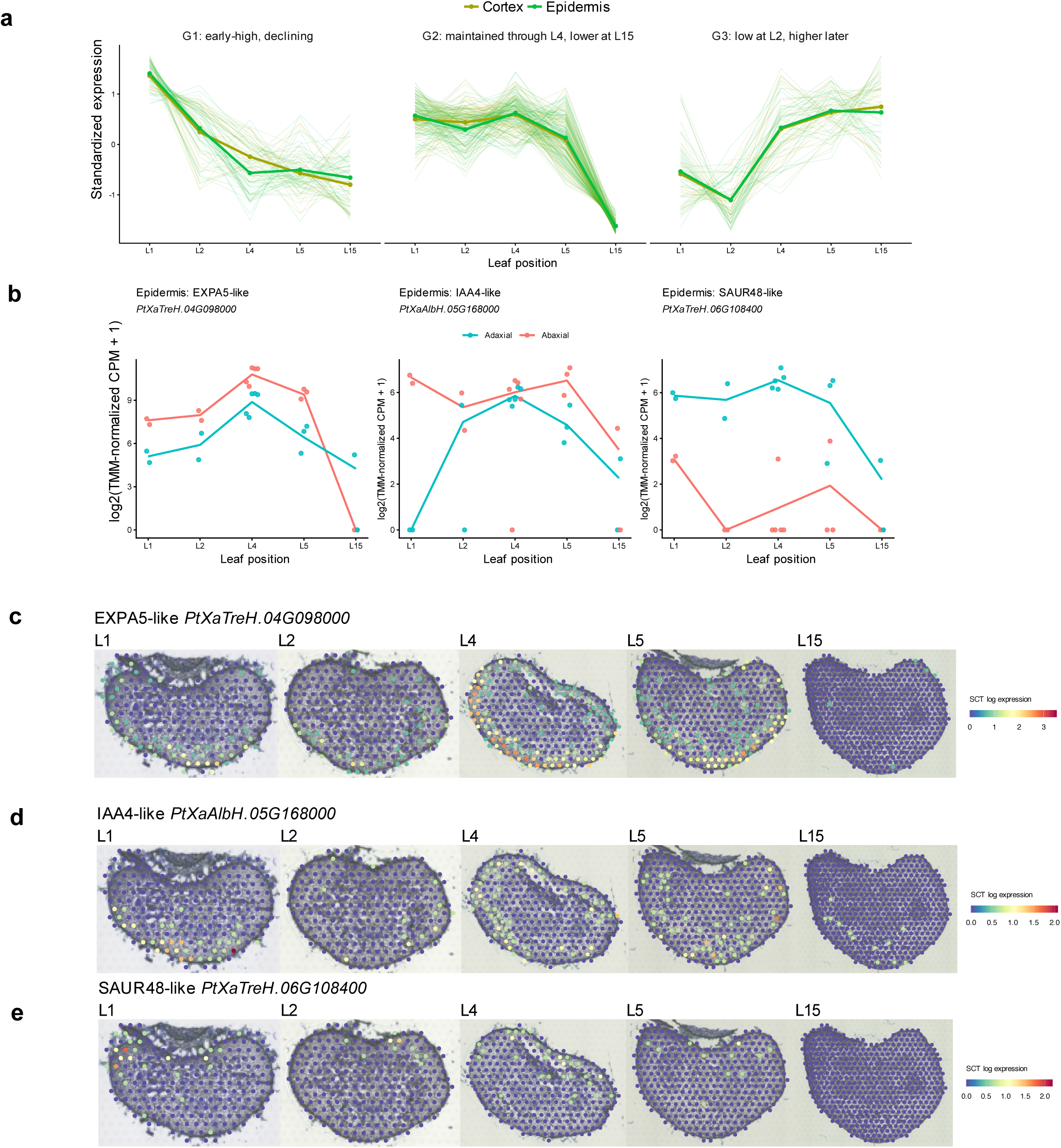
Developmental expression of petiole adaxial–abaxial DEGs. (a) Three groups of standardized expression profiles in epidermis and cortex. Thin curves show core profiles; thick curves show tissue-specific means. (b) Profiles of EXPA5-like, IAA4-like and SAUR48-like genes. (c–e) Corresponding spatial expression across five leaf positions. Colors indicate SCT log-normalized expression.

